# Transcranial *in vivo* detection of amyloid-beta at single plaque resolution with large-field multifocal illumination fluorescence microscopy

**DOI:** 10.1101/2020.02.01.929844

**Authors:** Ruiqing Ni, Zhenyue Chen, Gloria Shi, Alessia Villois, Quanyu Zhou, Paolo Arosio, Roger M. Nitsch, K. Peter R. Nilsson, Jan Klohs, Daniel Razansky

## Abstract

The abnormal deposition of beta-amyloid proteins in the brain is one of the major histopathological hallmarks of Alzheimer’s disease. Currently available intravital microscopy techniques for high-resolution plaque visualization commonly involve highly invasive procedures and are limited to a small field-of-view within the rodent brain. Here, we report the transcranial detection of amyloid-beta deposits at the whole brain scale with 20 μm resolution in APP/PS1 and arcAβ mouse models of Alzheimer’s disease amyloidosis using a large-field multifocal (LMI) fluorescence microscopy technique. Highly sensitive and specific detection of amyloid-beta deposits at a single plaque level in APP/PS1 and arcAβ mice was facilitated using luminescent conjugated oligothiophene HS-169. Immunohistochemical staining with HS-169, anti-Aβ antibody 6E10, and conformation antibodies OC (fibrillar) of brain tissue sections further showed that HS-169 resolved compact parenchymal and vessel-associated amyloid deposits. The novel imaging platform offers new prospects for *in vivo* studies into Alzheimer’s disease mechanisms in animal models as well as longitudinal monitoring of therapeutic responses at a single plaque level.

## Introduction

The abnormal accumulation and spread of amyloid-beta (Aβ) deposits are implicated to play a central role in the pathogenesis of Alzheimer’s disease (AD). Different conformations, aggregation states of Aβ, including Aβ monomer, oligomer, fibrillar Aβ and plaque, elicit different responses such as synaptotoxicity, neurotoxicity and inflammation (1–3). In a clinical setting, using positron emission tomography (PET) imaging with amyloid tracers such as ^11^C-PIB (4), ^18^F-florbetaben (5) and ^18^F-florbetapir (6), higher cortical fibrillar Aβ loads were reported in patients with AD and mild cognitive impairment compared to healthy controls (7). As a result, amyloid PET imaging has been established as an imaging biomarker for early and differential diagnosis of AD (8).

*In vivo* Aβ detection and its longitudinal monitoring in mouse models of AD amyloidosis has provided insights on the disease mechanisms and treatment effects. The detection has been possible at macroscopic level by using 3D microPET imaging with ^11^C-PIB (9), ^18^F-florbetapir (10), ^18^F-florbetaben (11), ^18^F-flutemetamol, ^11^C-AZD2184 (12), and ^125^I-labeled-antibody (13, 14). Alternatively, optical detection of Aβ deposits can be done with *ex vivo* optical projection tomography (15) as well as various *in vivo* planar fluorescence approaches (16) and assisted with various near-infrared contrast agents, such as AOI987 (17), CRANAD-2/-3 (18, 19), luminescent conjugated oligothiophenes (LCOs) (20, 21), or in 3D via fluorescence molecular tomography employing AOI987 (22). Imaging of Aβ at a higher (mesoscopic) resolution has also been demonstrated using ^19^F- and ^1^H-magnetic resonance imaging (23) or using dual modalities near-infrared magnetic resonance imaging (24), as well as optoacoustic tomography (25).

To this end, *in vivo* imaging of Aβ deposits at a single plaque resolution in an intact mouse brain may enable the understanding of growth dynamics of amyloid deposits at their earliest onset and evaluation of Aβ clearing therapies. As the diameter of Aβ plaques in murine models of amyloidosis range between 8-120 μm, the resolution of aforementioned macroscopic and mesoscopic imaging methods is insufficient for single Aβ deposit detection (26–28). Optoacoustic (29) and multiphoton microscopy techniques using methoxy-X04, BTA-1 and PIB (30–33) have been shown capable of monitoring Aβ with μm resolution. However, these techniques provide a limited field-of-view while commonly involving cranial opening, which may affect brain physiology.

We devised a large-field multi-focal illumination (LMI) fluorescence microscopy method that provides a unique combination between an extended 20 × 20 mm field-of-view as well as high spatial (~20 μm) and temporal (10 Hz) resolutions (34, 35). In the present study, we demonstrated whole brain mapping of Aβ deposits at single plaque resolution in APP/PS1 (26) and arcAβ (36) mouse models of AD cerebral amyloidosis mediated by HS-169 LCOs (37). Both strains are commonly used in AD research, but differ in their Aβ pathologies. The *in vivo* LMI imaging results were validated by *ex vivo* LMI imaging and immunohistochemistry using HS-169 with anti-Aβ antibody 6E10 and anti-amyloid fibrillar conformation antibody (OC) on mouse brain sections.

## Methods

### In vitro binding between amyloid probes HS-169 and recombinant Aβ_42_ fibrils

Recombinant Aβ_42_ monomers were expressed and produced by *E.coli* as described previously (38, 39). Amyloid fibrils were formed by incubating a solution of 2 μM Aβ_42_ in phosphate buffer (PBS, pH 8.0). The aggregation process was monitored by a quantitative fluorescence assay based on the Thioflavin T (ThT) dye (39). Fluorescence imaging *in vitro* of 2 μl of 30 μM HS-169, 30 μM HS169 + 1 μM Aβ_42_ fibril, and 1 μM Aβ_42_ fibril mixtures were performed at 0 and 30 minutes after mixing.

### Animal model

Two APP/PS1 mice (26) overexpressing the human APP695 transgene containing the Swedish (K670N/M671L) and PSEN1 containing an L166P mutations under the control of Th1 promoter and two age-matched non-transgenic littermates of both sexes (16 months-of-age) were used. In APP/PS1 mice, human Aβ42 is preferentially generated over Aβ40, but levels of both increase with age. In the brain, the Aβ_42_/Aβ_Flmi_ decreases with the onset of amyloid deposition (26). Amyloid plaque deposition starts at approximately six weeks of age in the neocortex. Deposits appear in the hippocampus at about three to four months, and in the striatum, thalamus, and brainstem at four to five months. Cognitive impairment have been reported to start at seven months of age (40). In addition, two arcAβ mice overexpressing the human APP695 transgene containing the Swedish (K670N/M671L) and Arctic (E693G) mutations under the control of prion protein promoter and two age-matched non-transgenic littermates of both sexes were used (24 months-of-age) (41). By six months of age, arcAβ mice develop amyloid pathology affecting both the brain parenchyma and vasculature. In the parenchyma, amyloid pathology starts as intracellular punctate Aβ deposits in the cortex and hippocampus. Plaques are abundant in these areas by 9 to 15 months (42). Severe cerebral amyloid angiopathy is also present by 9 to 15 months of age, with dense Aβ aggregates accumulating in the walls of blood vessels (42). Cerebral amyloid angiopathy leads to hypoperfusion, impaired vascular reactivity, decreased vessel density, blood-brain barrier impairment and occurrence of cerebral microbleeds (36, 43–46). Animals were housed in ventilated cages inside a temperature-controlled room, under a 12-hour dark/light cycle. Pelleted food (3437PXL15, CARGILL) and water were provided *ad-libitum*. All experiments were performed in accordance with the Swiss Federal Act on Animal Protection and were approved by the Cantonal Veterinary Office Zurich (permit number: ZH082/18).

### *In vivo* and *ex vivo* LMI fluorescence imaging of Aβ deposits in mice

Our recently developed LMI fluorescence imaging method based on a beam-splitting grating and an acousto-optic deflector synchronized with a high speed camera was employed for this study (**Fig. 1A**) (35). Briefly, a high-repetition pulsed Q-switched, diode end-pumped Nd:YAG laser (model: IS8II-E, EdgeWave, Germany) operating at 532 nm wavelength was used for the excitation. The laser beam was first scanned by the acousto-optic deflector AOD (AA Opto-Electronic, France) at 1 kHz and then guided into a customized beam-splitting grating (Holoeye GmbH, Germany) to generate 21×21 mini-beams. The mini-beams were relayed by 4f system and then focused onto the sample (mouse brain) to generate multiple foci, as shown in **Fig. 1A**. After passing through the dichroic mirror, the emitted fluorescence signal was collected and focused onto the sensor plane of a high-speed camera (PCO AG, Germany).

**Figure 1.**
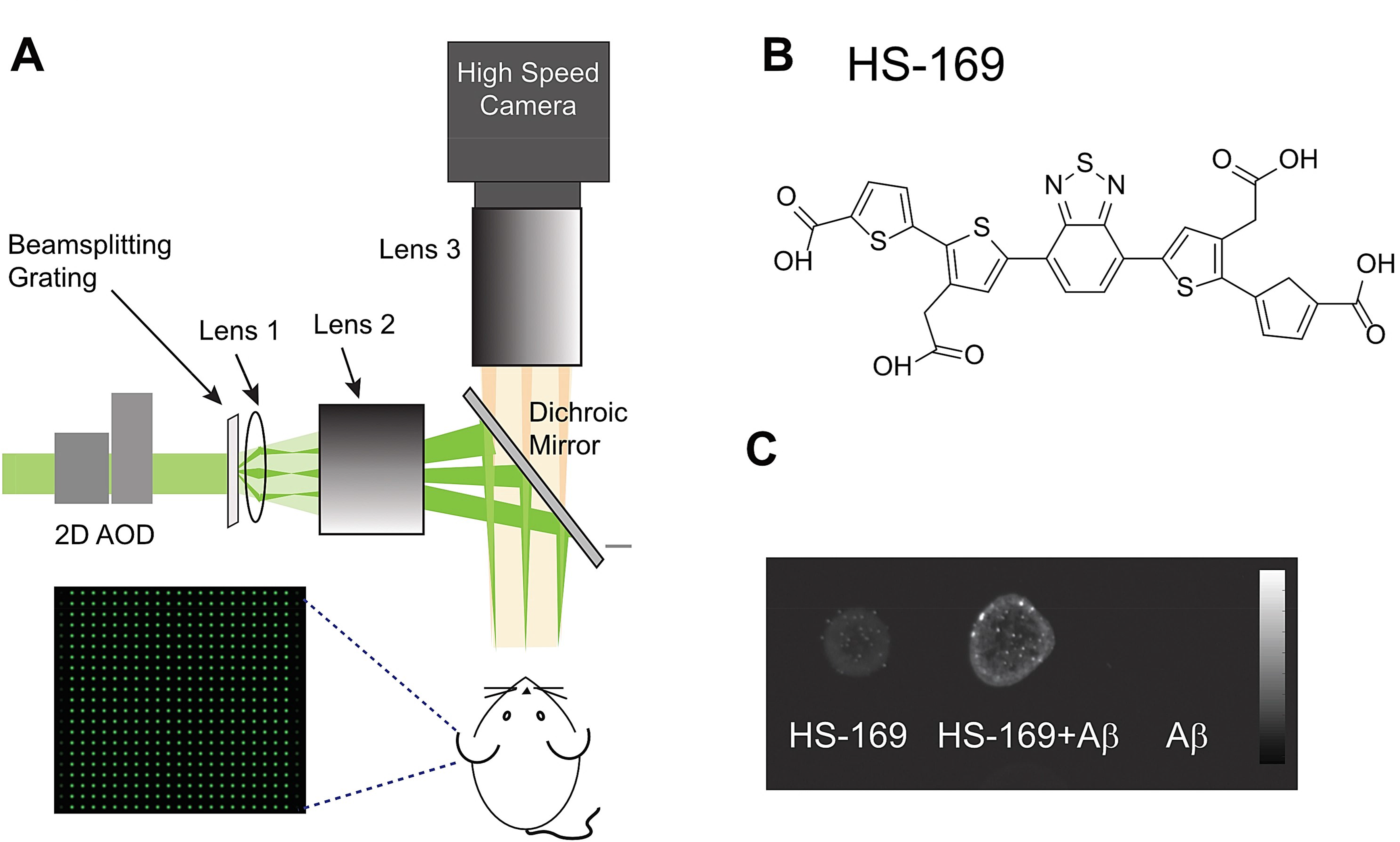
Large-field multifocal illumination (LMI) fluorescence microscopy system. (A) Schematic diagram of the set up consisting of an acousto-optic deflector (AOD), beam-splitting grating, focusing assembly, and a high-speed fluorescence camera; insert: illumination grid when a 21×21 beam-splitting grating is employed; (B) Chemical structure of amyloid imaging probe HS-169; (C) Increase of fluorescence intensity of HS-169 (30 μM) in aggregated Aβ_42_ fibril (1 μM) at 30 minutes after incubation.

The fluorescence LCO probe HS-169 (**Fig. 1B**) was synthesized as described previously (37). Mice were anesthetized with isoflurane (4 % v/v for induction and 1.5 % v/v during experiments) in 20 % O_2_ at a flow rate of ~0.5 l/min. Before imaging each mouse was positioned onto the imaging stage, the scalp was removed to reduce light scattering while the skull was kept intact. For APP/PS1, arcAβ mice and non-transgenic littermates, an *i.v.* tail-vein bolus injection of 0.4 mg/kg HS-169 solution in 0.1 M PBS (pH 7.4) was administered.

Next we compared the difference between the LMI imaging and conventional wide-field imaging. Brain sections from one APP/PS1, one arcAβ and one non-transgenic littermate mouse were imaged *ex vivo* after the *in vivo* imaging sessions using the aforementioned setup. The mice were sacrificed under deep anesthesia (ketamine/xylazine/acepromazine maleate (75/10/2 mg/kg body weight, i.p. bolus injection)) without prior perfusion. Horizontal brain slices of 3-mm thickness were cut in a brain matrix using a razor blade. Sections were placed on object holders wrapped with black tape.

To validate the *in vivo* and *ex vivo* signal, the other APP/PS1, arcAβ mouse and non-transgenic littermate were perfused under ketamine/xylazine/acepromazine maleate anesthesia (75/10/2 mg/kg body weight, i.p. bolus injection) with 0.1 M PBS (pH 7.4) and decapitated. The skulls were removed. The brains were then imaged using LMI imaging and afterwards post-fixed in 4 % paraformaldehyde in 0.1M PBS (pH 7.4) for 1 day and stored in 0.1 M PBS (pH 7.4) at 4 °C.

### Image reconstruction and data analysis

Reconstruction of the LMI images was performed based on the saved raw data from the CCD camera. Firstly, local maxima in each scanning frame were identified and the excited fluorescence signals were extracted with their centroids and intensity stored for later image stitching. Since the illumination grid was well defined with equal intervals between adjacent spots, this prior information facilitated signal extraction while suppressing noise. Correction for non-uniform beam intensity profile was then performed across each frame before combining the image values of local maxima to form the complete high-resolution image. Note that laser pulse energy fluctuations and other aberrations caused by optical components may necessitate additional corrections to obtain fine-tuned image quality and more uniform responsivity, however no such corrections were applied here. Fluorescence intensity (F.I.) in regions of interest (ROIs) were quantified for the dataset of *in vivo* (**Fig. 2)** and *ex vivo* imaging (**Fig. 3**). Full width at half maximum (FWHM) at x- and y-axis were used for plaque size analysis. Contrast-to-noise ratio (CNR) was calculated using the *in vivo* LMI imaging and conventional wide-field fluorescence microscope data acquired at 100 minutes after injections into APP/PS1, arcAβ and non-transgenic littermate mice.

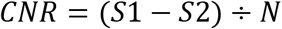

Where *S* is fluorescence intensity in the ROI; and *N* is the standard deviation from a region in the background.

**Figure 2.**
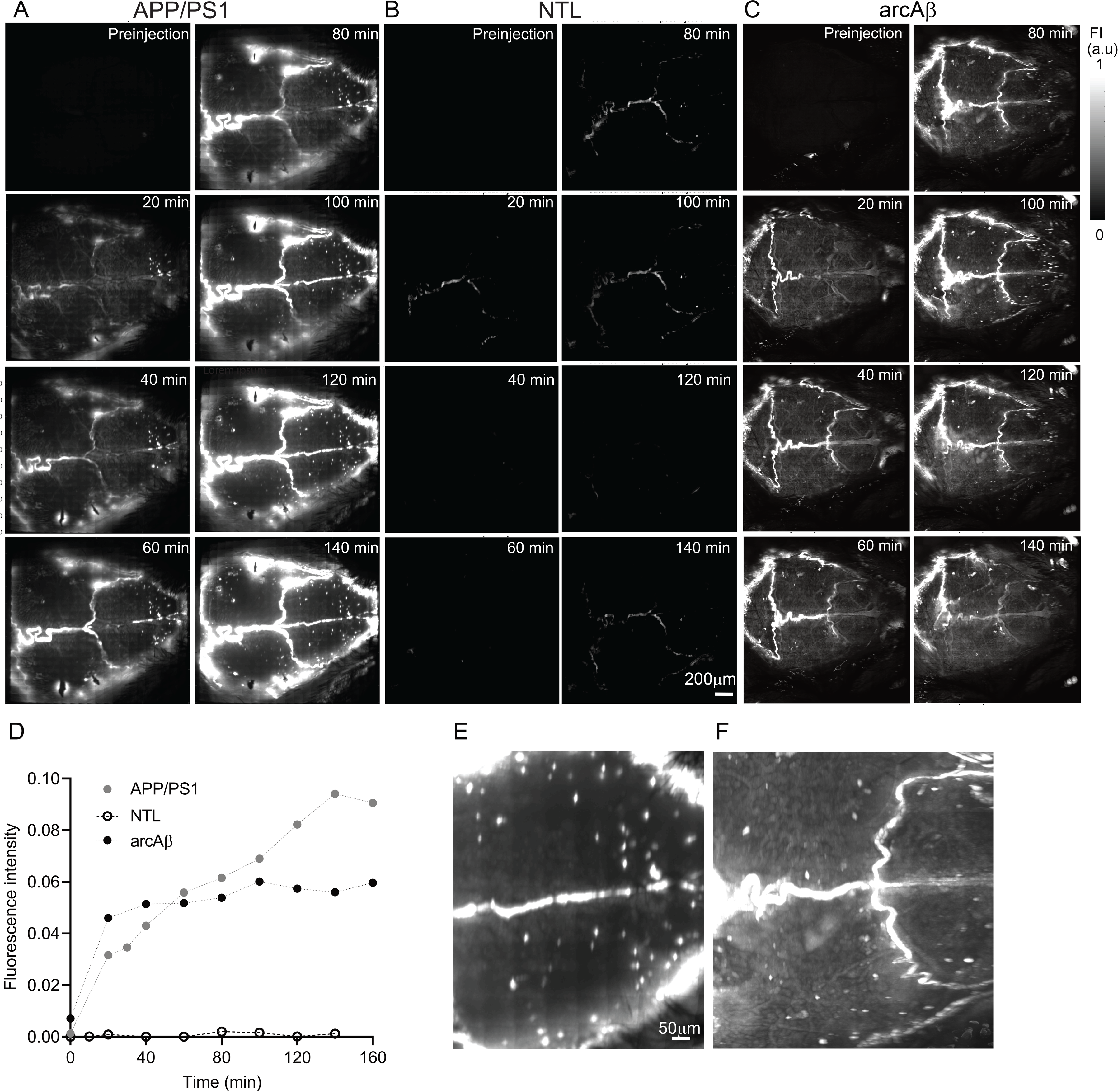
*In vivo* large-field multifocal illumination (LMI) imaging of amyloid-beta deposits in APP/PS1 and arcAβ mouse brain. (A) APP/PS1, (B) non-transgenic littermate (NTL) and (C) arcAβ mouse at pre-injection and followed up to 140 minutes after i.v. injection of HS169 through tail vein. Scale bar = 200 μm. Fluorescence intensity (FI) scale = 0-1 (a.u.); (D) Fluorescence intensity normalized to pre-injection values as a function of time after probe injection in APP/PS1, arcAβ and NTL mouse brains; (E) Zoom-in view of skull area over the cortex of an APP/PS1 at 140 minutes and (F) an arcAβ mouse at 100 minutes after injection. Scale bar = 50 μm;

**Figure 3.**
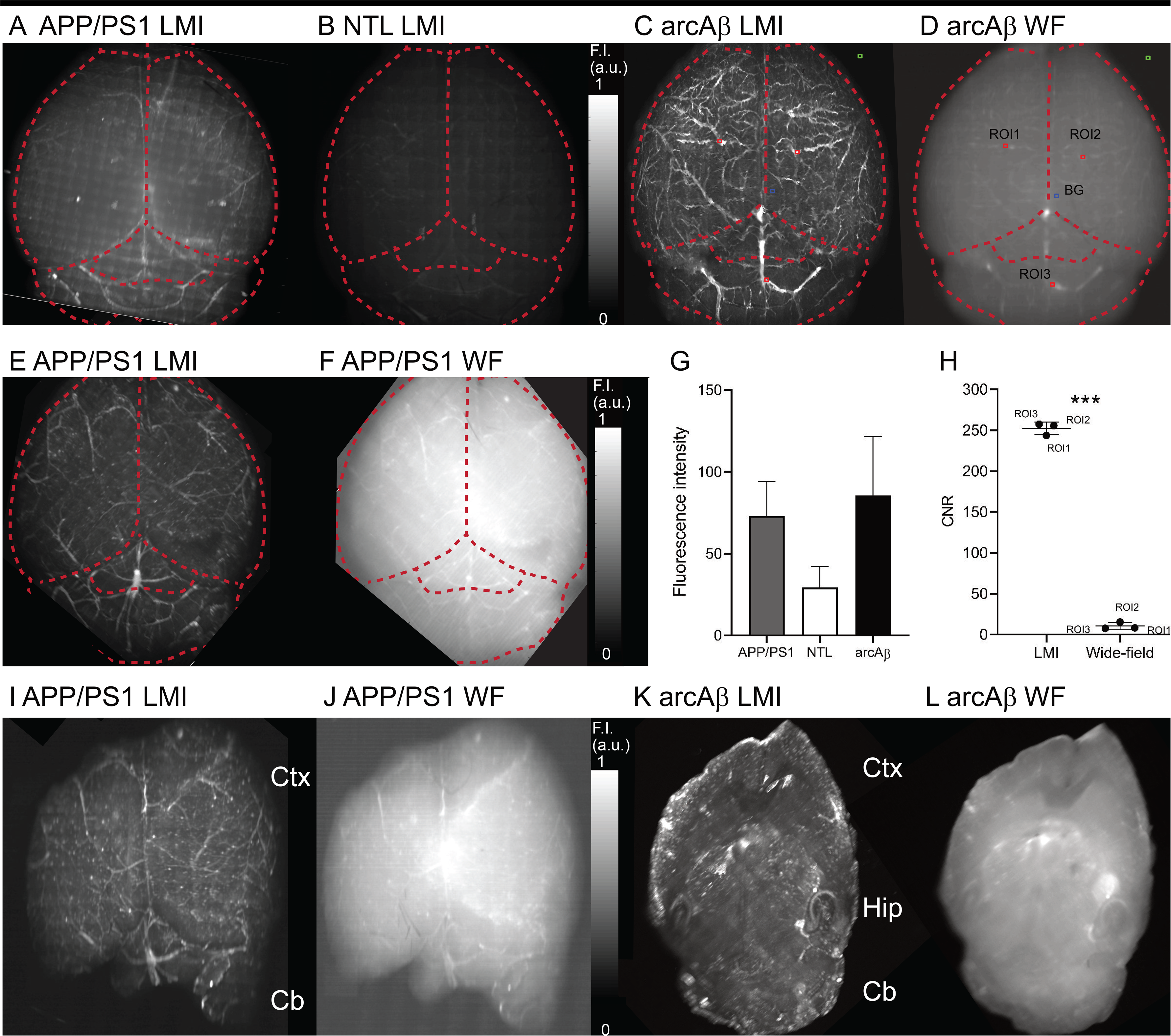
*Ex vivo* large-field multifocal illumination (LMI) imaging compared to wide-field fluorescence microscopy of amyloid-beta deposits in APP/PS1 and arcAβ mouse brain. LMI imaging *ex vivo* showed higher signal in the whole brain from (A) one APP/PS1 and (C) one arcAβ mouse compared to (B) one non-transgenic littermate; Higher contrast to noise is further achieved by using (C, E) LMI imaging than (D, F) wide-field (WF) fluorescence imaging in the *ex vivo* whole brain of APP/PS1 and arcAβ mouse; (G) Quantification of signal of *ex vivo* LMI imaging in APP/PS1, arcAβ mouse and non-transgenic littermate (A-C); (H) contrast to noise ratio (CNR) of e*x vivo* LMI imaging and WF imaging in one arcAβ mouse; ***p<0.001; Region of interest (ROIs, red squares) and background (BG, blue square), noise (green square) indicated in C, D; (I, J) *ex vivo* LMI imaging and WF imaging of brain slice from APP/PS1 mouse; (K, L) *ex vivo* whole brain LMI imaging in arcAβ (Fluorescence intensity scale = 0-1 (a.u.)). Scale bar = 50 μm. Ctx: cortex; Hip: hippocampus; Cb: cerebellum.

### Immunohistochemical staining and confocal microscopy

Histology and immunohistochemical investigations were performed on the mouse brain sections after PBS perfusion and *ex vivo* LMI imaging. Brain hemispheres were embedded in paraffin following routine procedures and were cut in 5 μm horizontal sections. For immunohistochemical staining, 6E10, HS-169, and fibrillar conformation anti-amyloid antibodies OC (47) stainings were performed following protocol described earlier with nuclei counterstained by 4′,6-diamidino-2-phenylindole (DAPI) (44) (Details in **Supplementary Table 1**). Histochemical staining using Hematoxylin & Eosin were performed for structural information and detecting of abnormalities in the brain. The whole brain slices of 6E10 and Alexa488, Cyanine3, OC were imaged at ×20 magnification using Pannoramic 250 (3D HISTECH, Hungary) at the ScopeM core imaging facility of the ETH for quality control of the autofluorescence and antibody specificity.

To further assess the co-localization of different channels, confocal images of arcAβ, APP/PS1 mice and non-transgenic littermates were further obtained at ×10, × 63 magnification in the cortex, hippocampus areas, and ×20 magnification for the whole brain slices using a Leica SP8 confocal microscope (Leica Microsystems GmbH, Germany) at the ScopeM. Sequential images were obtained by using 405 nm, 488 nm, 561 nm lines respectively. Identical resolution settings were used for the Z stacks (n = 15). Full width at half maximum (FWHM) at x- and y-axis were used for plaque size analysis using ×10 cortex confocal images for both HS-169 and 6E10 channels. The Allen brain atlas was used for anatomical reference (48).

### Statistics

Paired two-tail *student t* test was used (Graphpad Prism) for comparing values between LMI imaging and wide-field imaging. All data are present as mean ± standard deviation. Significance was set at * *p* < 0.05.

## Results

### *In vitro* LCO binding in aggregated Aβ_42_ fibrils

Very weak fluorescence was detected shortly after co-incubation of HS-169 with Aβ_42_ fibrils. The signals were significantly increasing over time reaching 3756.7: 1552.2: 725.8 (a.u.) for HS169+Aβ: HS-169: Aβ at 30 minutes post co-incubation (**Fig. 1C**).

### *In vivo* LMI imaging using LCO exhibited stronger fluorescence signal from the brain in APP/PS1 and arcAβ mice compared to NTLs

After *i.v*. injection of HS-169 through the mouse tail vein, fluorescence intensity increase was observed in the mouse brain indicating that the probe was passing the blood-brain barrier. In general, the HS-169 fluorescence intensity increased over time in the brain from both APP/PS1 and arcAβ mice, although exhibiting different kinetics. In APP/PS1 mice the signal increase was slower peaking at 140 minutes (**Fig. 2A**) while in arcAβ mice (**Fig. 2C**) the signal increased faster in the early phase peaking at 100 minutes. The highest signal intensity was observed at 100-140 minutes post-injection indicating specific binding rather than probe wash-in. In comparison, HS-169 fluorescence intensity remained low in the brain from non-transgenic littermates (**Fig. 2B**). At 120 minutes, fluorescence signal integrated over the cortex was 663 times higher in the brain from APP/PS1 mouse and 463 times higher in arcAβ mouse in comparison with that in non-transgenic littermate (**Fig. 2D).** Bright signal spots appeared on LMI fluorescence microscopy images, in both the brains from APP/PS1 and arcAβ mice, suggestive of single Aβ deposits. Size analysis showed that the spots detected in the brain from APP/PS1 and arcAβ mice are of mean 31 μm, range 15-150 μm, resembling typical plaque sizes (**Fig. 2E-F**). It should be noted that accuracy of the size analysis is influenced by the fluorescence blooming effect as well as the difficulty to separate two closely neighboring signal sources in the fluorescence microscope. Intriguingly, these single signal spots were observed in APP/PS1 and arcAβ mouse in the area overlaying the cortex after 40minutes and remained visible until the end of data acquisition (140-160 minutes), substantiating the capacity to visualize individual plaques (**Fig. 2A-C**).

### *Ex vivo* LMI imaging using LCO HS-169 showed stronger cortical signal in APP/PS1, arcAβ mice as compared to non-transgenic littermates

*Ex vivo* imaging was performed on mouse brain slices as well as on PBS-perfused whole brains. *Ex vivo* LMI imaging revealed higher levels of cortical HS-169 accumulation in APP/PS1 and arcAβ mice compared to non-transgenic littermates (**Fig. 3A-C)**. Greater visibility and thus detection sensitivity of Aβ plaques were observed by LMI imaging as compared to wide-field (WF) imaging both in whole brains (**Fig. 3C-F**) as well as in individual horizontal brain sections (**Fig. 3I-L**). The fluorescence signal integrated over the cortical areas was approximately 2.5 times higher in the APP/PS1 and arcAβ mice compared to non-transgenic mice (**Fig. 3F)**. The contrast-to-noise ratio (CNR) was approximately 24 times higher in LMI imaging compared to wide-field images (t test, p = 0.0003) (**Fig. 3G,** ROI indicated in **Fig. 3C, D**). Compared to APP/PS1 mice, the HS-169 signal in the brain vasculature became more apparent in arcAβ mice.

### Regional distribution of plaques and difference in probe binding

Immunohistochemical and histological staining were performed on horizontal brain tissue sections from APP/PS1, arcAβ mice and non-transgenic littermates using HS-169, 6E10, OC and DAPI (**Fig. 4**). In APP/PS1, the Aβ plaques distributed mainly in the parenchymal regions (pronounced in the cortex and hippocampus) (**Fig. 4A, B**), and the size of plaques identified with 6E10 (mean 27 μm, range 4-117 μm) was larger as compared to HS-169 (mean 19 μm, range 2-82 μm) (**Fig. 4M**). In arcAβ mice, both parenchymal (pronounced in the cortex, hippocampus and thalamus) and vascular deposits are detected. The plaques in arcAβ mice are less diffuse compared to that in APP/PS1 mice. Plaques identified by using 6E10 (mean 40 μm, range 7-131 μm) are also higher in diameter than HS-169 mean 27 μm, range 2-117 μm) in arcAβ mice. OC staining for fibrillar Aβ showed good overlap with HS-169 in the cortical slices from both APP/PS1 and arcAβ mice (**Fig. 4N-P**). The lack of 6E10 and HS-169 signal in cortex of the non-transgenic littermate indicated highly specific binding of HS-169 to Aβ which absent in these brains (**Fig. 4R**).

**Figure 4.**
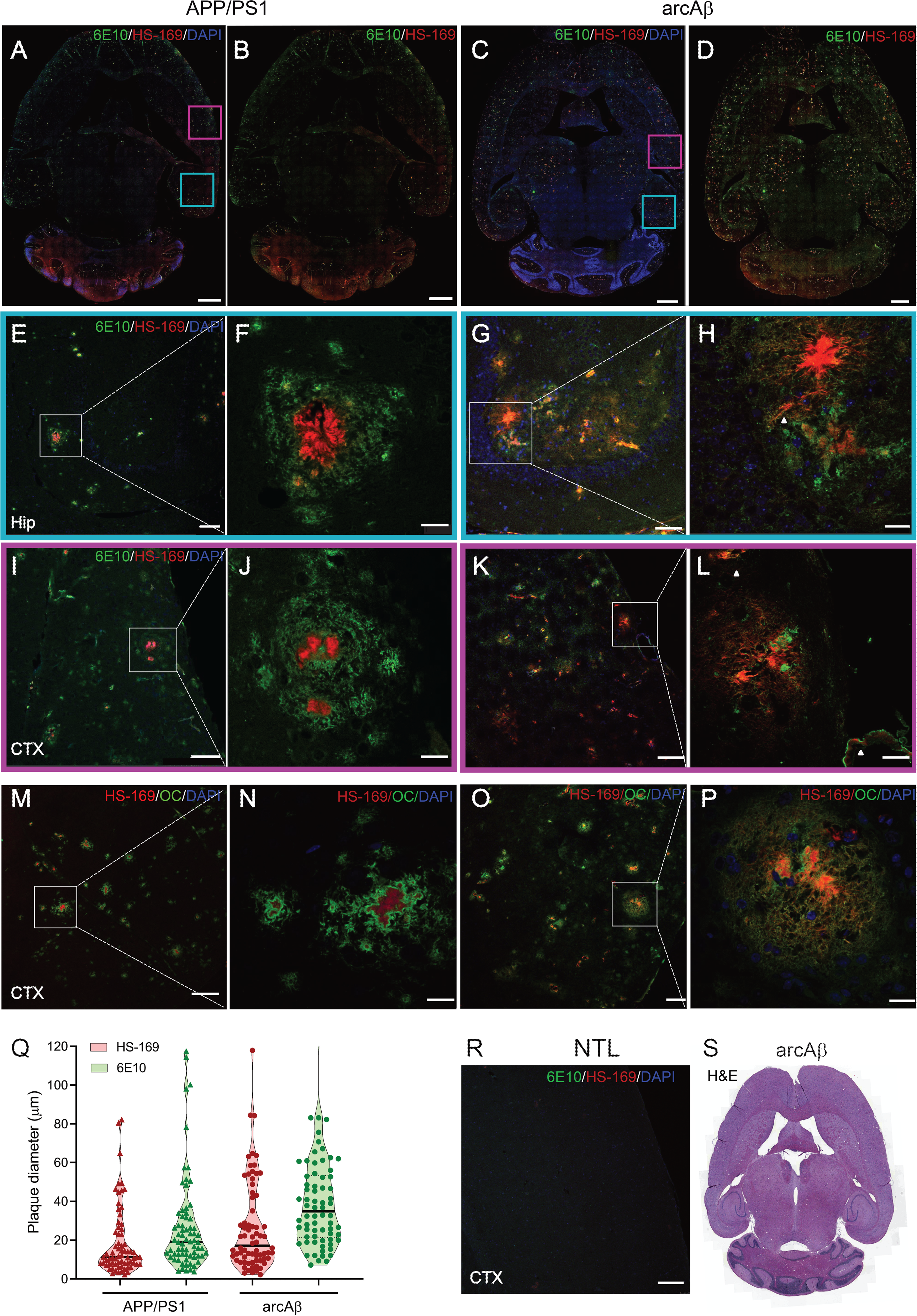
Staining for Aβ deposition in APP/PS1 and arcAβ mouse brain tissue sections. (A, B) Confocal imaging in horizontal whole brain sections from APP/PS1 mouse; and (C, D) from arcAβ mouse. DAPI (blue), Alexa488-6E10 (green), HS-169 (red); Zoom in of (E-H) hippocampus (blue square); (I-L) cortex (magenta square) in (A, C) respectively; demonstrating co-localization of 6E10 and HS-169 to amyloid-beta plaque in APP/PS1 and arcAβ mouse brain; (M) Better overlapping between HS-169 and 6E10 in small compact amyloid deposits; (N-P) Confocal imaging in cortex from APP/PS1 and arcAβ mice, OC (yellow), Alexa488-6E10 (green), HS-169 (red), DAPI (blue); (Q) Size analysis shows the plaque detected using 6E10 and HS-169 in the cortex from APP/PS1 and arcAβ mice; (R) No positive 6E10 or HS-169 signal in the cortex section from non-negative control mice; (S) Hematoxylin & Eosin staining on horizontal brain section from one arcAβ mouse. White arrowhead = cerebral amyloid angiopathy; Scale bar =1 mm (A-D); 100 μm (E, G, I, K, O, R); 20 μm (F, H, J, L, N, P).

## Discussion

Developing tools for non-invasive detection of Aβ deposits at high-resolution is important for understanding disease mechanism and translational development of Aβ-targeted disease-modifying therapies. Here, we demonstrated a novel *in vivo* LMI imaging approach to detect brain Aβ deposits at single plaque resolution with a large field-of-view covering the entire cortex in APP/PS1 and arcAβ mouse models.

In LMI imaging the out-of-focus light can be effectively rejected, due to the inherent advantage of laser beam scanning. Thus the proposed method enables minimally invasive imaging *in vivo* without employing craniotomy, a powerful advantage especially when it comes to longitudinal imaging of disease progression and treatment monitoring in aging mouse models of AD amyloidosis. By employing an image reconstruction algorithm that first extracts signals from small foci then superimposes them to form a high resolution image, the LMI imaging approach further enables high resolving power at a single plague level across the entire mouse cortex. Compared to laser scanning confocal microscopy, LMI imaging is a highly parallelized technique employing hundreds of illumination foci, thus enabling fast imaging speed which is crucial for mitigating motion artifacts in *in vivo* studies. In addition, the LMI imaging approach is optimally suited for imaging large objects up to a centimeter scale, which is not attainable with multi-photon microscopy methods that are further hindered by the lack of optimal labels with large absorption cross section and longer absorption/emission wavelengths (49, 50). Due to the small numerical aperture of the mini-beams, a large depth of focus has been further achieved, which makes the system suitable for imaging highly curved surfaces such as the mouse head. Yet, due to the lack of the optical sectioning ability, the current LMI imaging implementation can only obtain 2D information from the mouse brain.

*In vivo* imaging of Aβ plaques with other imaging modalities, such as PET, confronts the challenge of an inadequate spatial resolution to distinguish individual plaques (11, 12, 51–53). Moreover, mouse brain imaging by PET is limited by a complicated experimental setting, short half-life of positron-emitting nuclei (17), spillover as well as severe partial volume effects (54). MRI has been reported to detect Aβ plaques from APP/PS1 mice (23, 55, 56), albeit its low detection sensitivity is generally inadequate for detecting amyloid plaque due to its insufficient contrast against the surrounding tissues (56).

LMI amyloid imaging *in vivo* and *ex vivo* using HS-169 in APP/PS1 and arcAβ mice showed patterns generally fitting the known Aβ distribution in these mouse lines and was confirmed by immunohistochemistry. In APP/PS1 mice, abundant Aβ deposits were detected in the cortex and hippocampus (26) by using immunohistochemical staining, while arcAβ mouse showed higher presence of cortical, hippocampal and thalamic plaques and higher load of cerebral amyloid angiopathy (36, 43). The higher load of amyloid in the vessel wall might explain the faster kinetics of probe accumulation in arcAβ mice compared to APP/PS1 mice. Size analysis (28) of the Aβ deposits detected *in vivo* by HS-169and *ex vivo* by HS-169 and 6E10 revealed comparable values to what was previously reported in both APP/PS1 (28, 57) and arcAβ mouse brains (36). We observed that HS-169 stained the core of Aβ deposits *ex vivo*, thus the area stained by HS-169 is smaller than that by 6E10. In addition, HS-169 stained less proportion of 6E10 plaque in APP/PS1 compared to in arcAβ mouse. This is probably due to the more fibrillar composition and compact plaque structure in the arcAβ mouse (as observed in 6E10 and OC staining, **Fig. 4H, P**) as compared to the larger and more diffuse plaque types in APP/PS1 (**Fig. 4F, J**). In the smaller compact Aβ deposits (**Fig. 4M**), more overlapping between 6E10 and HS-169 was observed. As chemical probes such as HS-169 chiefly detect the beta-sheet structure, which is richer in the compact fibrillar Aβ, higher similarity is expected between HS-169 and 6E10 in the arcAβ than in APP/PS1 mouse brain. Different amyloid composition in mouse models of amyloidosis has been reported in previous studies by using immunohistochemical staining (58–60). In human brain, different binding property of various probes to amyloid deposits in familial AD patients with various mutation were also reported (61). Mean diameters for HS-169 were estimated to be 19 μm and 27 μm for APP/PS1 and arcAβ mice respectively, which corresponds well with spot sizes of 10-30 μm measured with LMI imaging *in vivo*. However, it indicates also that plaques sizes are underestimated (stronger in arcAβ mice) with LMI imaging using HS-169 and might limit the capability to follow individual plaque growth. Thus, in the future novel probes may be developed and tested, which show better correspondence with true Aβ plaque size (27).

APP/PS1 is a widely used animal model for AD mechanistic study (62–66). The single Aβ deposits detection described in the current work is directly applicable to study Aβ plaque clearance (67–70), and to monitor antibody treatment effect in the mouse models (9, 62, 71, 72), which can currently only be achieved by craniotomy and *ex vivo* techniques. Serial *in vivo* LMI imaging of the same mice would thus significantly reduce the number of animals required for studies monitoring Aβ accumulation, compared to cross-sectional analysis with immunohistochemistry. In addition, the suggested LMI imaging method may potentially provide a new platform for detecting tau deposits (21, 73) and other protein aggregates of beta-sheet structures (74) *in vivo* in animal models using chemical probes. Note that we have previously shown efficacy of the single plaque detection framework in arcAβ mouse model (36, 45). Yet, further studies on younger mice (to determine detection thresholds), and other mouse strains of amyloidosis such as APPswe, APPswe/PS1dE9 mice(75) are essential for corroborating the presently available results. Additional limitations include confounding factors such as the skull thickness of the mice, and vascular abnormality in the transgenic mouse lines (especially of old age), which may have influenced the detected fluorescence intensity levels.

In conclusion, we demonstrated *in vivo* high-resolution whole brain Aβ imaging by LMI imaging without skull opening in APP/PS1 and arcAβ mouse model of AD amyloidosis. The new imaging platform offers new prospects for *in vivo* studies into AD-related disease mechanisms in animal models as well as longitudinal monitoring of therapeutics targeting Aβ.

## Availability of data and material

The datasets generated and/or analyzed during the current study are available in the repository DOI zenodo 10.5281/zenodo.3564424.

## Author Contributions

RN, ZC, JK, DR conceived and designed the study; RN, ZC, GS, QZ, AV performed the experiments; KPRN provided HS-169, RN, ZC, GS analyzed the data; RN, ZC, JK, DR interpreted the results; RN, ZC, JK, DR wrote the paper; all coauthors contributed constructively to the manuscript.

## Acknowledgement

The authors acknowledge technical support from Dr. Joachim Hohls of the Scientific Center for Optical and Electron Microscopy (ScopeM) of ETH Zurich, Ms Marie Renault at Institute for Biomedical Engineering of ETH Zurich, and Ms Priyanka Ravikumar at Institute for Regenerative Medicine of University of Zurich for technical help.

## References

1. Walsh DM, et al. (2002) Naturally secreted oligomers of amyloid beta protein potently inhibit hippocampal long-term potentiation in vivo. Nature 416(6880):535–539.

2. Zott B, et al. (2019) A vicious cycle of beta amyloid-dependent neuronal hyperactivation. Science 365(6453):559–565.

3. De Strooper B & Karran E (2016) The Cellular Phase of Alzheimer’s Disease. Cell 164(4):603–615.

4. Klunk WE, et al. (2005) Binding of the positron emission tomography tracer Pittsburgh compound-B reflects the amount of amyloid-beta in Alzheimer’s disease brain but not in transgenic mouse brain. The Journal of neuroscience: the official journal of the Society for Neuroscience 25(46):10598–10606.

5. Barthel H, et al. (2011) Cerebral amyloid-β PET with florbetaben (18F) in patients with Alzheimer’s disease and healthy controls: a multicentre phase 2 diagnostic study. The Lancet Neurology 10(5):424–435.

6. Fleisher AS, et al. (2011) Using Positron Emission Tomography and Florbetapir F 18 to Image Cortical Amyloid in Patients With Mild Cognitive Impairment or Dementia Due to Alzheimer Disease. Archives of Neurology 68(11):1404–1411.

7. Villemagne VL, Dore V, Burnham SC, Masters CL, & Rowe CC (2018) Imaging tau and amyloid-beta proteinopathies in Alzheimer disease and other conditions. Nat Rev Neurol 14(4):225–236.

8. Jack CR, Jr., et al. (2018) NIA-AA Research Framework: Toward a biological definition of Alzheimer’s disease. Alzheimer’s & dementia: the journal of the Alzheimer’s Association 14(4):535–562.

9. Snellman A, et al. (2017) Applicability of [(11)C]PIB micro-PET imaging for in vivo follow-up of anti-amyloid treatment effects in APP23 mouse model. Neurobiol Aging 57:84–94.

10. Poisnel G, et al. (2012) PET imaging with [18F]AV-45 in an APP/PS1-21 murine model of amyloid plaque deposition. Neurobiology of Aging 33(11):2561–2571.

11. Yousefi BH, et al. (2015) FIBT versus florbetaben and PiB: a preclinical comparison study with amyloid-PET in transgenic mice. EJNMMI Res 5:20.

12. Rodriguez-Vieitez E, et al. (2015) Astrocytosis precedes amyloid plaque deposition in Alzheimer APPswe transgenic mouse brain: a correlative positron emission tomography and in vitro imaging study. European journal of nuclear medicine and molecular imaging 42(7):1119–1132 (cofirst author).

13. Sehlin D, et al. (2016) Antibody-based PET imaging of amyloid beta in mouse models of Alzheimer’s disease. Nature communications 7:10759.

14. Syvanen S, et al. (2017) A bispecific Tribody PET radioligand for visualization of amyloid-beta protofibrils - a new concept for neuroimaging. NeuroImage 148:55–63.

15. Nguyen D, et al. (2019) Supervised learning to quantify amyloidosis in whole brains of an Alzheimer’s disease mouse model acquired with optical projection tomography. Biomed Opt Express 10(6):3041–3060.

16. Pansieri J, et al. (2019) Ultraviolet–visible–near-infrared optical properties of amyloid fibrils shed light on amyloidogenesis. Nature Photonics 13(7):473–479.

17. Hintersteiner M, et al. (2005) In vivo detection of amyloid-beta deposits by near-infrared imaging using an oxazine-derivative probe. Nat Biotechnol 23(5):577–583.

18. Ran C, et al. (2009) Design, synthesis, and testing of difluoroboron-derivatized curcumins as near-infrared probes for in vivo detection of amyloid-beta deposits. J Am Chem Soc 131(42):15257–15261.

19. Zhang X, et al. (2015) Near-infrared fluorescence molecular imaging of amyloid beta species and monitoring therapy in animal models of Alzheimer’s disease. Proceedings of the National Academy of Sciences 112(31):9734.

20. Åslund A, et al. (2009) Novel Pentameric Thiophene Derivatives for in Vitro and in Vivo Optical Imaging of a Plethora of Protein Aggregates in Cerebral Amyloidoses. ACS chemical biology 4(8):673–684.

21. Calvo-Rodriguez M, et al. (2019) In vivo detection of tau fibrils and amyloid beta aggregates with luminescent conjugated oligothiophenes and multiphoton microscopy. Acta Neuropathol Commun 7(1):171.

22. Hyde D, et al. (2009) Hybrid FMT-CT imaging of amyloid-beta plaques in a murine Alzheimer’s disease model. NeuroImage 44(4):1304–1311.

23. Higuchi M, et al. (2005) 19F and 1H MRI detection of amyloid beta plaques in vivo. Nat Neurosci 8(4):527–533.

24. Li Y, et al. (2018) Dual-Modal NIR-Fluorophore Conjugated Magnetic Nanoparticle for Imaging Amyloid-beta Species In Vivo. Small 14(28):e1800901.

25. Ni R, Vaas M, Rudin M, & Klohs J (2018) Quantification of amyloid deposits and oxygen extraction fraction in the brain with multispectral optoacoustic imaging in arcAbeta; mouse model of Alzheimer’s disease. SPIE BiOS 10494:6.

26. Radde R, et al. (2006) Abeta42-driven cerebral amyloidosis in transgenic mice reveals early and robust pathology. EMBO Rep 7(9):940–946.

27. Whitesell JD, et al. (2019) Whole brain imaging reveals distinct spatial patterns of amyloid beta deposition in three mouse models of Alzheimer’s disease. The Journal of comparative neurology 527(13):2122–2145.

28. Jahrling N, et al. (2015) Cerebral beta-Amyloidosis in Mice Investigated by Ultramicroscopy. PloS one 10(5):e0125418.

29. Hu S, Yan P, Maslov K, Lee J-M, & Wang LV (2009) Intravital imaging of amyloid plaques in a transgenic mouse model using optical-resolution photoacoustic microscopy. Opt. Lett. 34(24):3899–3901.

30. Hefendehl JK, et al. (2011) Long-term in vivo imaging of beta-amyloid plaque appearance and growth in a mouse model of cerebral beta-amyloidosis. The Journal of neuroscience: the official journal of the Society for Neuroscience 31(2):624–629.

31. Meyer-Luehmann M, et al. (2008) Rapid appearance and local toxicity of amyloid-beta plaques in a mouse model of Alzheimer’s disease. Nature 451(7179):720–724.

32. Klunk WE, et al. (2002) Imaging Aβ Plaques in Living Transgenic Mice with Multiphoton Microscopy and Methoxy-X04, a Systemically Administered Congo Red Derivative. Journal of Neuropathology & Experimental Neurology 61(9):797–805.

33. Bacskai BJ, et al. (2003) Four-dimensional multiphoton imaging of brain entry, amyloid binding, and clearance of an amyloid-β ligand in transgenic mice. Proceedings of the National Academy of Sciences 100(21):12462–12467.

34. Chen Z, et al. (2018) Multifocal structured illumination fluorescence microscopy with large field-of-view and high spatio-temporal resolution (SPIE).

35. Chen Z, et al. (High-Speed Large-Field Multifocal Illumination Fluorescence Microscopy. Laser & Photonics Reviews n/a(n/a):1900070.

36. Merlini M, Meyer EP, Ulmann-Schuler A, & Nitsch RM (2011) Vascular beta-amyloid and early astrocyte alterations impair cerebrovascular function and cerebral metabolism in transgenic arcAbeta mice. Acta neuropathologica 122(3):293–311.

37. Shirani H, et al. (2015) A Palette of Fluorescent Thiophene-Based Ligands for the Identification of Protein Aggregates. Chemistry 21(43):15133–15137.

38. Arosio P, et al. (2016) Kinetic analysis reveals the diversity of microscopic mechanisms through which molecular chaperones suppress amyloid formation. Nature communications 7:10948.

39. Cohen SIA, et al. (2015) A molecular chaperone breaks the catalytic cycle that generates toxic Aβ oligomers. Nature Structural & Molecular Biology 22:207.

40. Serneels L, et al. (2009) gamma-Secretase heterogeneity in the Aph1 subunit: relevance for Alzheimer’s disease. Science 324(5927):639–642.

41. Knobloch M, Konietzko U, Krebs DC, & Nitsch RM (2007) Intracellular Abeta and cognitive deficits precede beta-amyloid deposition in transgenic arcAbeta mice. Neurobiol Aging 28(9):1297–1306.

42. Knobloch M, Konietzko U, Krebs D, & Nitsch R (2007) Intracellular Abeta and cognitive deficits precede beta-amyloid deposition in transgenic arcAbeta mice. Neurobiol Aging 28(9):1297–1306.

43. Klohs J, et al. (2013) Longitudinal Assessment of Amyloid Pathology in Transgenic ArcAbeta Mice Using Multi-Parametric Magnetic Resonance Imaging. PloS one 8(6):e66097.

44. Ni R, et al. (2019) fMRI Reveals Mitigation of Cerebrovascular Dysfunction by Bradykinin Receptors 1 and 2 Inhibitor Noscapine in a Mouse Model of Cerebral Amyloidosis. Front aging neurosci 11:27–27.

45. Klohs J, et al. (2016) Quantitative assessment of microvasculopathy in arcAβ mice with USPIO-enhanced gradient echo MRI. J Cereb Blood Flow Metab 36(9):1614–1624.

46. Ielacqua GD SF, Füchtemeier M, Xandry J, Rudin M, Klohs J. (2016) Magnetic Resonance Q Mapping Reveals a Decrease in Microvessel Density in the arcAβ Mouse Model of Cerebral Amyloidosis. Front Aging Neurosci 7:241.

47. Kayed R, et al. (2007) Fibril specific, conformation dependent antibodies recognize a generic epitope common to amyloid fibrils and fibrillar oligomers that is absent in prefibrillar oligomers. Mol Neurodegener 2:18.

48. Jones AR, Overly CC, & Sunkin SM (2009) The Allen Brain Atlas: 5 years and beyond. Nature reviews. Neuroscience 10(11):821–828.

49. Kim HM & Cho BR (2015) Small-molecule two-photon probes for bioimaging applications. Chem Rev 115(11):5014–5055.

50. Jun YW, et al. (2019) Frontiers in Probing Alzheimer’s Disease Biomarkers with Fluorescent Small Molecules. ACS Central Science 5(2):209–217.

51. Ni R, Gillberg PG, Bergfors A, Marutle A, & Nordberg A (2013) Amyloid tracers detect multiple binding sites in Alzheimer’s disease brain tissue. Brain: a journal of neurology 136:2217–2227.

52. Brendel M, et al. (2015) Cross-Sectional Comparison of Small Animal [18F]-Florbetaben Amyloid-PET between Transgenic AD Mouse Models. PloS one 10(2):e0116678.

53. Snellman A, et al. (2013) Longitudinal amyloid imaging in mouse brain with 11C-PIB: comparison of APP23, Tg2576, and APPswe-PS1dE9 mouse models of Alzheimer disease. Journal of nuclear medicine: official publication, Society of Nuclear Medicine 54(8):1434–1441.

54. Brendel M, et al. (2014) Impact of partial volume effect correction on cerebral beta-amyloid imaging in APP-Swe mice using [(18)F]-florbetaben PET. NeuroImage 84:843–853.

55. Dhenain M, et al. (2009) Characterization of in vivo MRI detectable thalamic amyloid plaques from APP/PS1 mice. Neurobiology of Aging 30(1):41–53.

56. Jack CR, Jr., et al. (2005) In vivo magnetic resonance microimaging of individual amyloid plaques in Alzheimer’s transgenic mice. The Journal of neuroscience: the official journal of the Society for Neuroscience 25(43):10041–10048.

57. Yan P, et al. (2009) Characterizing the appearance and growth of amyloid plaques in APP/PS1 mice. The Journal of neuroscience: the official journal of the Society for Neuroscience 29(34):10706–10714.

58. Kulic L, et al. (2012) Early accumulation of intracellular fibrillar oligomers and late congophilic amyloid angiopathy in mice expressing the Osaka intra-Aβ APP mutation. Translational Psychiatry 2(11):e183–e183.

59. Kim HY, et al. (2015) EPPS rescues hippocampus-dependent cognitive deficits in APP/PS1 mice by disaggregation of amyloid-β oligomers and plaques. Nature communications 6(1):8997.

60. Willuweit A, et al. (2009) Early-Onset and Robust Amyloid Pathology in a New Homozygous Mouse Model of Alzheimer’s Disease. PloS one 4(11):e7931.

61. Ni R, et al. (2017) Amyloid tracers binding sites in autosomal dominant and sporadic Alzheimer’s disease. Alzheimer’s & dementia: the journal of the Alzheimer’s Association 13(4):419–430.

62. Tian X, et al. (2019) Multimodal Imaging of Amyloid Plaques: Fusion of the Single-Probe Mass Spectrometry Image and Fluorescence Microscopy Image. Anal Chem 91(20):12882–12889.

63. Wirths O, Weis J, Szczygielski J, Multhaup G, & Bayer TA (2006) Axonopathy in an APP/PS1 transgenic mouse model of Alzheimer’s disease. Acta neuropathologica 111(4):312–319.

64. van Groen T, Kiliaan AJ, & Kadish I (2006) Deposition of mouse amyloid β in human APP/PS1 double and single AD model transgenic mice. Neurobiology of Disease 23(3):653–662.

65. Trinchese F, et al. (2004) Progressive age-related development of Alzheimer-like pathology in APP/PS1 mice. Annals of neurology 55(6):801–814.

66. Ni R, Rudin M, & Klohs J (2018) Cortical hypoperfusion and reduced cerebral metabolic rate of oxygen in the arcAbeta mouse model of Alzheimer’s disease. Photoacoustics 10:38–47.

67. Pitschke M, Prior R, Haupt M, & Riesner D (1998) Detection of single amyloid beta-protein aggregates in the cerebrospinal fluid of Alzheimer’s patients by fluorescence correlation spectroscopy. Nat Med 4(7):832–834.

68. Burgold S, Filser S, Dorostkar MM, Schmidt B, & Herms J (2014) In vivo imaging reveals sigmoidal growth kinetic of β-amyloid plaques. Acta Neuropathologica Communications 2(1):30.

69. Yu R-J, et al. (2019) Single molecule sensing of amyloid-β aggregation by confined glass nanopores. Chemical Science.

70. Klohs J, Rudin M, Shimshek DR, & Beckmann N (2014) Imaging of cerebrovascular pathology in animal models of Alzheimer’s disease. Front Aging Neurosci 6:32.

71. van Dyck CH (2018) Anti-Amyloid-β Monoclonal Antibodies for Alzheimer’s Disease: Pitfalls and Promise. Biological psychiatry 83(4):311–319.

72. Maeda J, et al. (2007) Longitudinal, Quantitative Assessment of Amyloid, Neuroinflammation, and Anti-Amyloid Treatment in a Living Mouse Model of Alzheimer’s Disease Enabled by Positron Emission Tomography. The Journal of Neuroscience 27(41):10957.

73. Ni R, et al. (2018) Comparative in-vitro and in-vivo quantifications of pathological tau deposits and their association with neurodegeneration in tauopathy mouse models. Journal of nuclear medicine: official publication, Society of Nuclear Medicine 59(6):960–966.

74. Taylor JP, Hardy J, & Fischbeck KH (2002) Toxic proteins in neurodegenerative disease. Science 296(5575):1991–1995.

75. Garcia-Alloza M, et al. (2006) Characterization of amyloid deposition in the APPswe/PS1dE9 mouse model of Alzheimer disease. Neurobiology of Disease 24(3):516–524.

