## Supplementary Table 1 for "Transcranial *in vivo* detection of amyloid-beta at single plaque resolution with large-field multifocal illumination fluorescence microscopy"

**Supplementary Table 1.** List of primary antibodies and ligands used for immunohistochemistry

| **Antibody** | **Company** | **Cat. No.** | **Dilution** |
| --- | --- | --- | --- |
| Mouse anti-Ab-6E10 | Signet Lab | SIG-39320 | 1:5000 |
| DAPI | Sigma | D9542-10MG | 1:1000 |
| HS-169 |  |  | 5 μM |
| Donkey-anti-Rat Alexa 488 | Jackson | AB_2340686 | 1:400 |
| Mouse-anti-human monoclonal Alexa488-6E10 | Biolegend | SIG-39347 | 1:200 |
| Anti-amyloid fibrils OC Rabbit polyclonal | Merck | AB2286 | 1:200 |
| Goat-anti-Rabbit Cyanine3  Goat-anti-Rabbit Alexa488 | Invitrogen  Invitrogen | A10520  A11034 | 1:200  1:200 |
